# 1q21.3 amplification modulates therapy sensitivity in Triple Negative Breast Cancer

**DOI:** 10.1101/2020.04.29.066993

**Authors:** Ramakanth Chirravuri-Venkata, Dario Ghersi, Apar K. Ganti, Imayavaramban Lakshmanan, Sanjib Chaudary, Surinder K. Batra, Mohd Wasim Nasser

## Abstract

The contrast in therapy sensitivity and response across triple negative breast cancer (TNBC) patients suggest underlying genotypic heterogeneity. Using publicly available data, we found significant associations between DNA-level copy number alterations of 1q21.3 locus and therapy sensitivity. We show that in spite of their aggressive nature, 1q21.3 amplified tumors are more responsive to commonly used cytotoxic therapies, highlighting the relevance of 1q21.3 copy number status as a genetic marker for risk stratification, therapy selection and response.

## Introduction

Breast cancer (BRCa) is the most common malignancy among women in the US and is expected to affect 1 in 8 (13%) American women in their lifetime. Despite the availability of effective treatments and an advanced understanding of the disease biology, unaccountable differences in the therapy response across patients continue to exist. The triple negative (TNBC) subtype is the most aggressive form of BRCa, and is characterized by the lack of estrogen receptor (ER), progesterone receptor (PR), and HER2 receptor expression. It is generally assumed to carry a high likelihood of recurrence and rapid progression. Nevertheless, TNBC has long been considered a heterogeneous disease due to its diversity in genotype and therapy response as a subset of patients are very responsive to chemotherapy (1). Studies have demonstrated that ~40% of TNBC patients undergo pathological complete response (pCR) when treated with neoadjuvant chemotherapy and these patients have improved overall and event-free survival (2). This contrast in therapy response supports the possibility of underlying genetic modulators among TNBC patients. Thus, there is a great need for novel molecular markers for an evidence-based disease management of TNBC (3). Using publicly available data, we uncovered genotypic characteristics modulating the therapy outcomes in TNBC.

Genomic alterations are generally a consequence of replication stress (DRS) and xenobiotics like hydroxyurea and cisplatin may act as a trigger for such events (4,5). It is also known that some of the standard chemotherapeutic agents like doxorubicin can set off endoduplication events in cancer cells (6). However, several studies have also demonstrated the effect of copy number alterations on therapeutic efficacy and survival outcomes (7). Although there is increasing evidence suggesting the associations between xenobiotic efficacy (particularly cytotoxic agents) and genomic instability, it is not clear why particular alterations are much more common than others and how they modulate cellular dynamics. Furthermore, understanding the effect of cytogenetic abnormalities on therapy response and vice-versa would help understand the underlying biology of drug efficacy and help in the development of improved treatments.

As one of the largest chromosomes with ~3000 genes, chromosome 1 is often genetically altered. Some of these alterations are already linked to numerous medical conditions like cancer, diabetes and schizophrenia (8,9). Despite the wealth of information regarding these alterations, the context and time dependent changes in these alterations have been less explored in cancer. In particular, 1q amplification is a very common alteration in prostate (neuroendocrine and exocrine), breast, pancreas, and biliary system cancers. In BRCa, the alteration frequency is around 8-20% and the frequency is even higher (30-40%) in TNBC (10). Interestingly, 1q amplification is often termed as “regional amplification” due to its discontinuous amplification across the chromosomal arm or even within a particular locus of the arm (11).

Numerous studies have demonstrated the association between high expression of individual 1q genes like MUC1, S100 & SPRR family of genes with disease progression and therapy resistance (12). However, the major determinant of mRNA expression of most genes is inherently their copy number status (13). Therefore, elucidating the role played by DNA copy number changes in addition to mRNA expression can provide a more complete view. Although previous studies indicate that gain of DNA copy number of 1q genes is associated with poor prognostic outcomes, here we show that 1q21.3 amplification imparts selective sensitivity to cytotoxic therapies.

## Results

### Characterization of 1q21.3 amplified TNBC tumors using TCGA and METABRIC datasets

We used TCGA and METABRIC TNBC copy number alteration data to study the association between 1q21.3 amplification and survival of patients who received both chemotherapy and radiotherapy. Due to regional amplification in the locus, we used a set of three representative/proxy genes (S100A8, HORMAD1, and VPS72) for analysis. The genes were selected based on their chromosomal location, amplification incidence and the correlation between the mRNA levels and their DNA-level copy number (CN). Used in combination, the mean mRNA level of these genes is fairly associated with CN status (figure 1a). High incidence frequency of amplification events in the representative genes was observed in Basal (35%), Luminal (A = 28%, B = 22%) and HER2 (16%) molecular subtypes obtained by PAM50 + low claudin classifier (14) (figure 1b).

**Figure 1.**
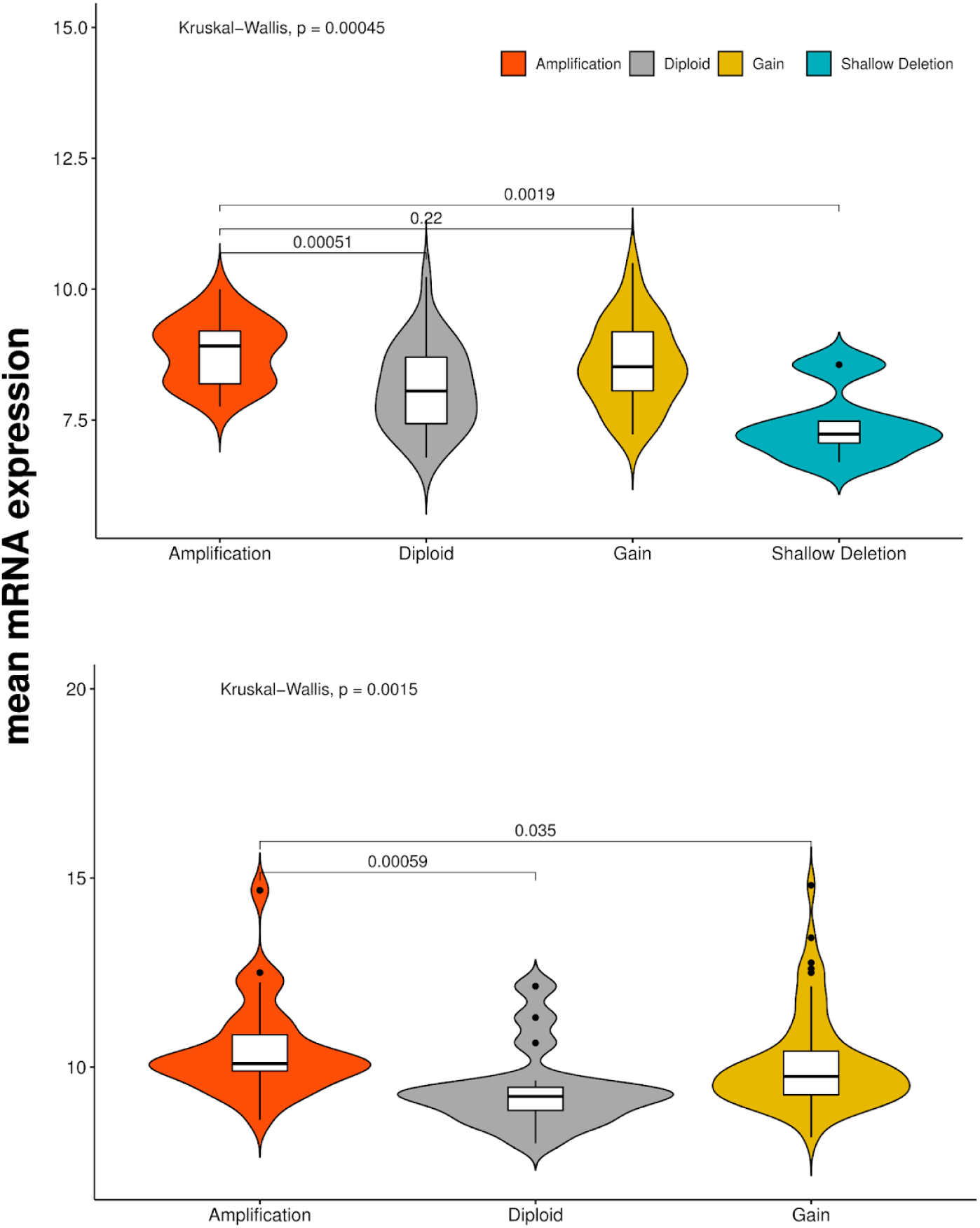
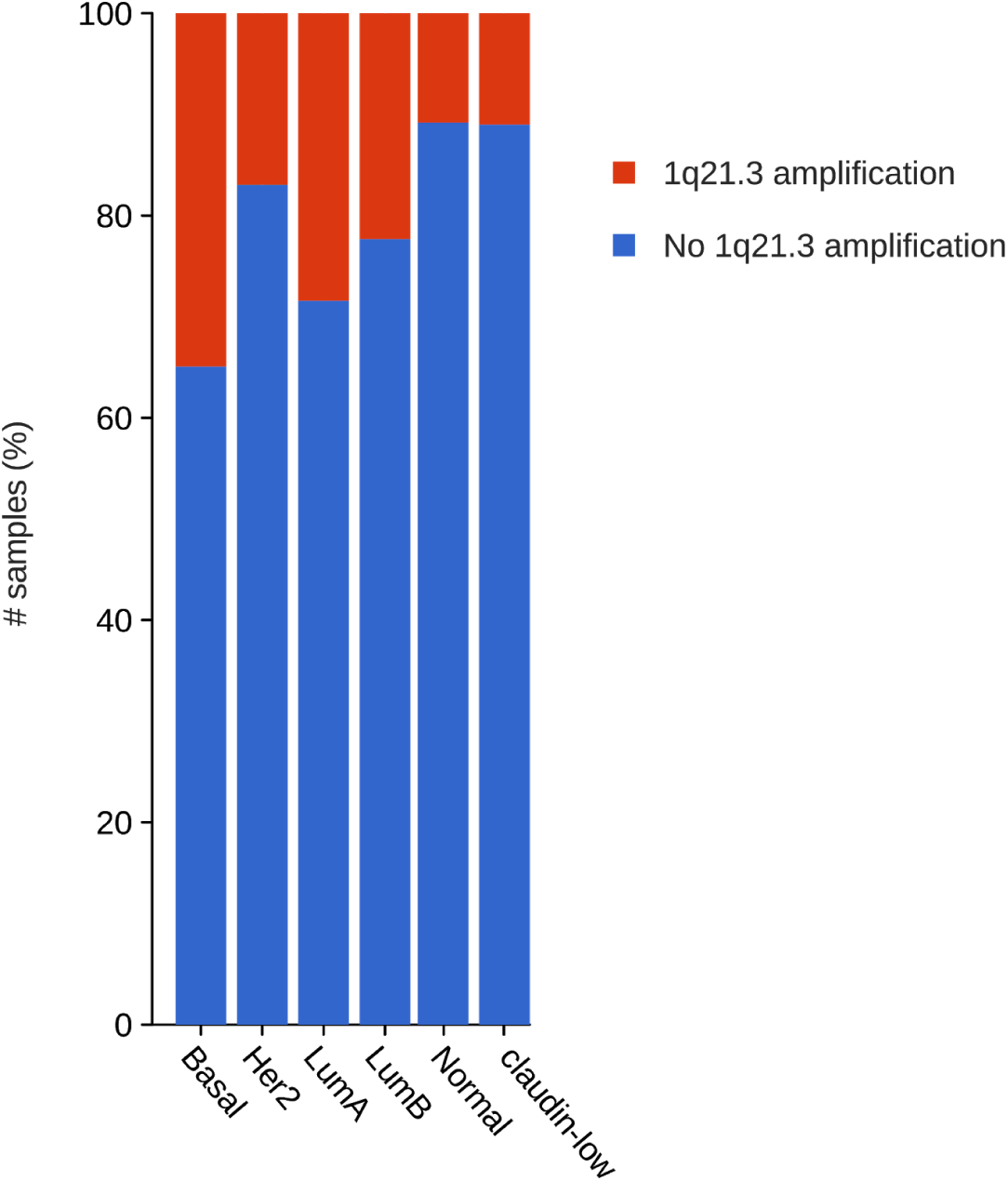
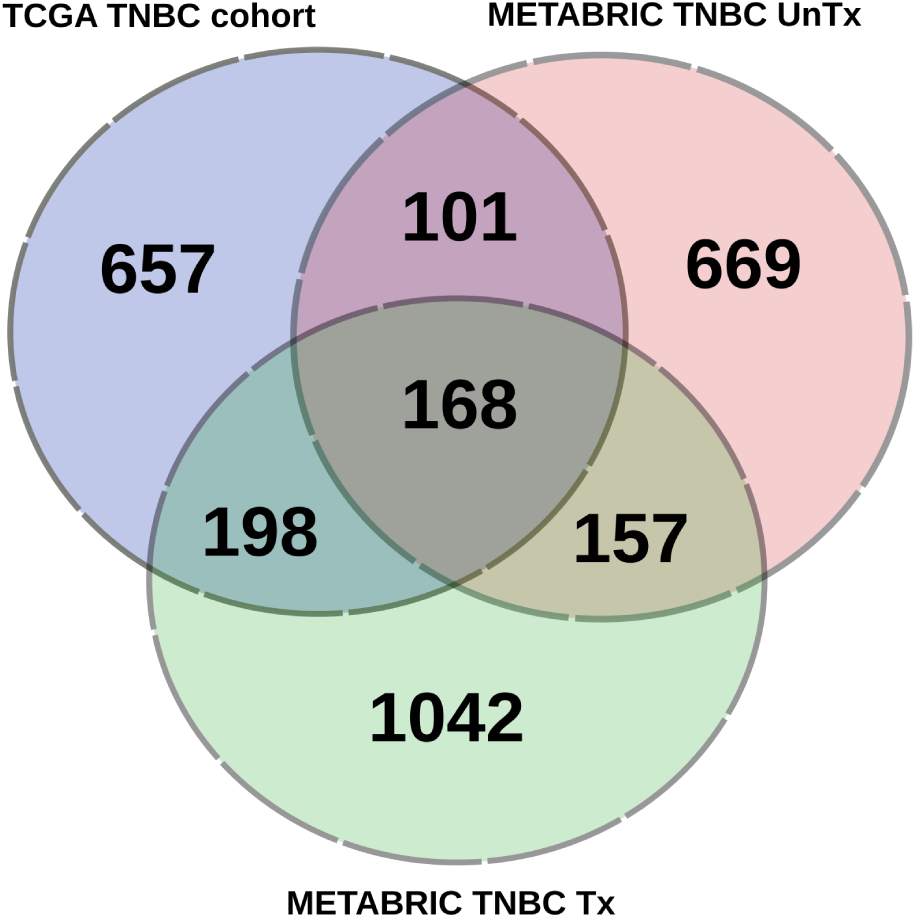
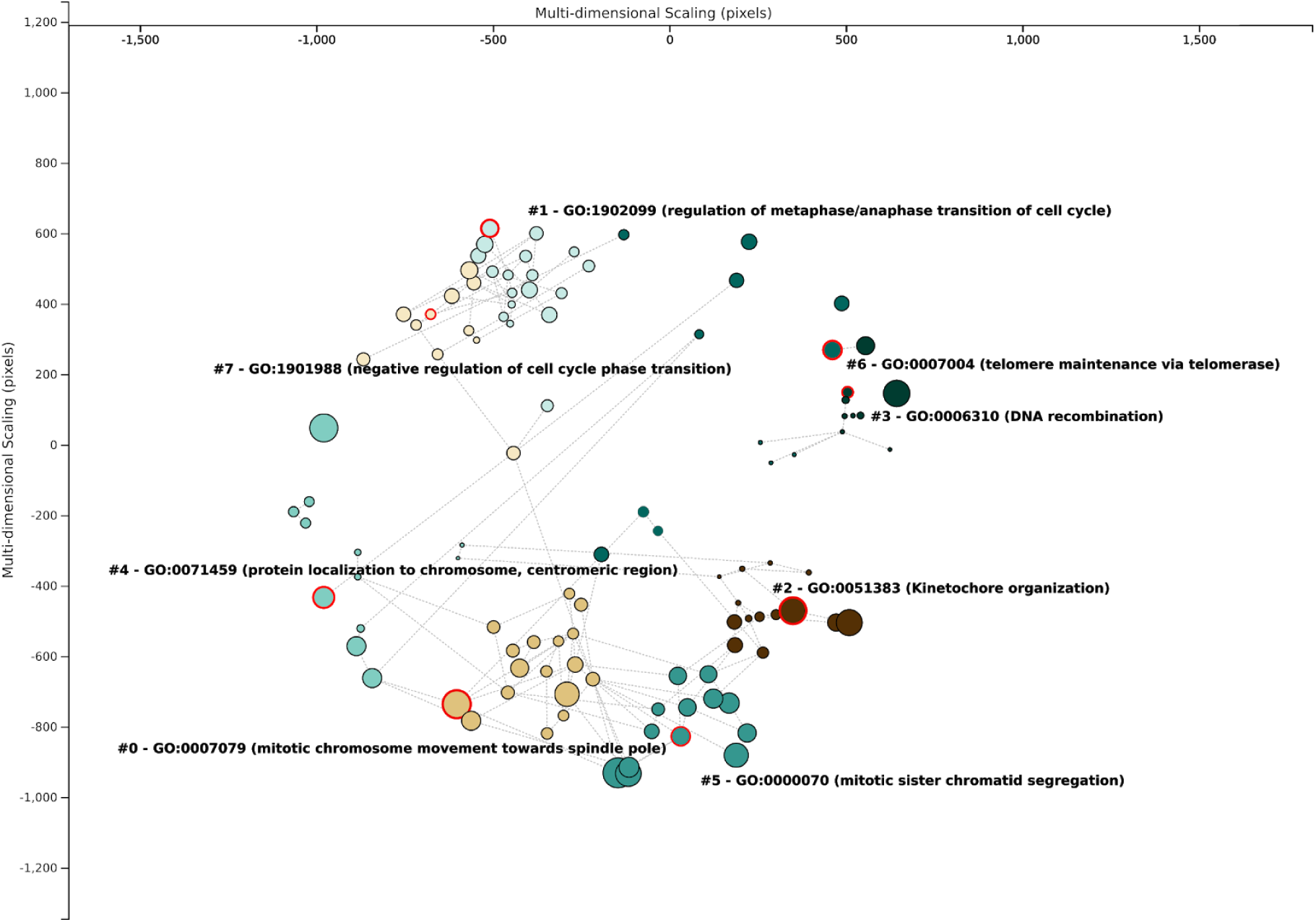
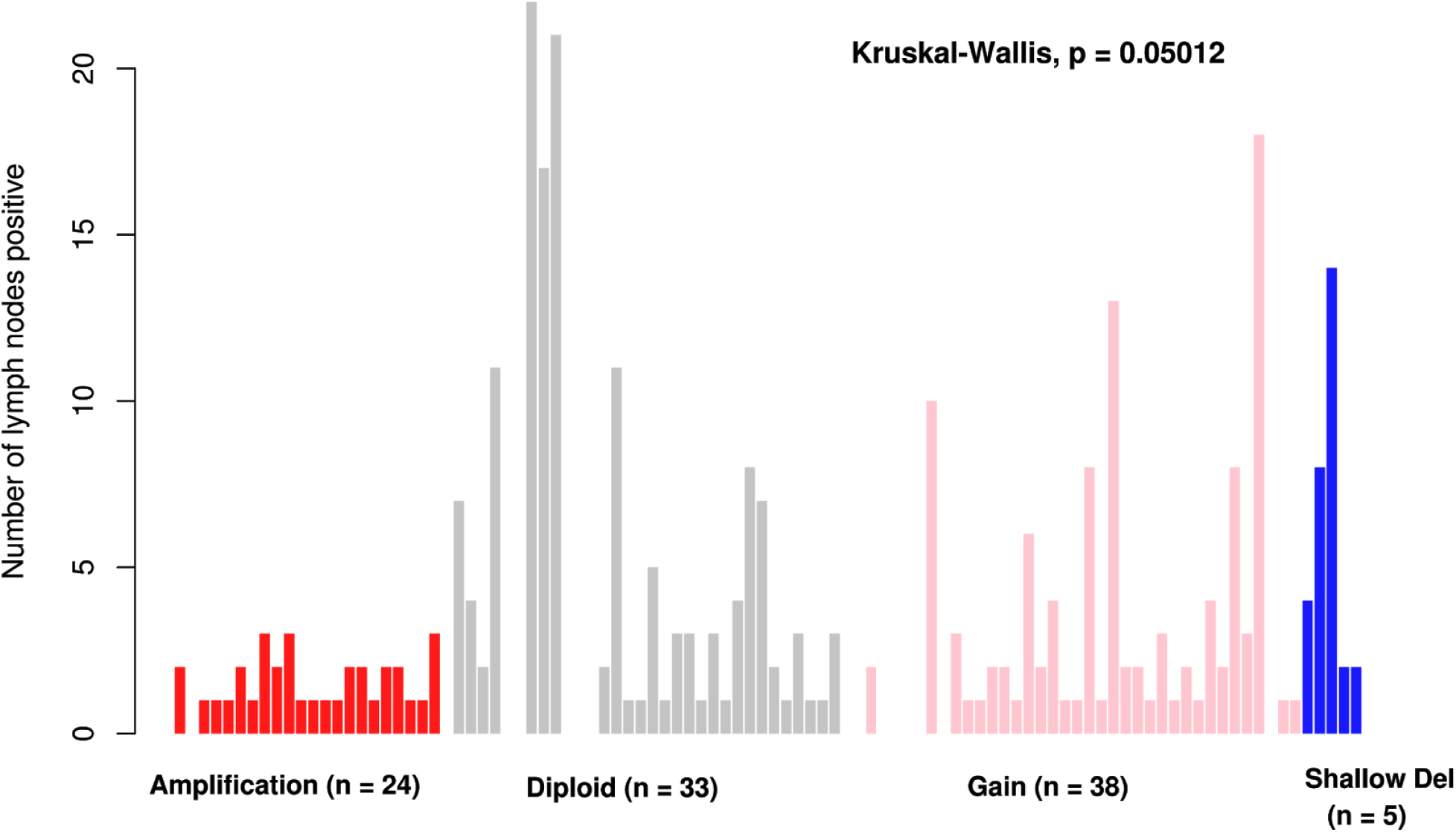
a) Plot illustrating the mean mRNA expression values of representative genes (S100A8, HORMAD1 & VPS72) with respect to their median copy number status (SD = Shallow deletion, D = Diploid, G = Gain & A = Amplification) using METABRIC treated (top; n = 100) and TCGA (bottom; n = 111) TNBC cohorts. Kruskal wallis p values are reported for multiple group comparisons and wilcoxon test p values are reported for pairwise comparisons. b) Incidence frequency of 1q21.3 amplification across molecular subtypes (PAM50 + low claudin classification) using METABRIC dataset (n = 2509). Highest incidence of 1q21.3 amplification is observed in basal subtype (35%) tumors followed by luminal (A = 28%, B = 22%) and HER2 (16%) subtypes. c) Venn diagram representing significantly over-expressed genes across 1q21.3 amplified samples of TCGA TNBC treatment-naive cohort (n = 112), METABRIC TNBC treated (n = 104) and untreated cohorts (n = 48). d) Bubble chart illustrating the Gene Ontology based biological processes of common over-expressed in 1q21.3 amplified samples across TCGA and METABRIC TNBC treated and untreated cohorts. e) Barplot illustrates the number of positive lymph node counts across METABRIC chemo-radiotherapy treated samples. Consensus (median) of copy number alterations of 1q21.3 representative genes was used to infer the type of alteration.

TNBC patients with amplification events in either of the representative genes exhibited significantly better overall and progression-free survival. Larger significant differences in survival were observed after stage homogenization (using only Stage II) (figure 2). Similar results were observed when the analysis was repeated using basal and low-claudin samples as defined by the PAM50 classification scheme (14). Pathway analysis using *gProfiler* on differentially over-expressed genes in amplified samples showed overrepresentation of mitotic pathways (15) (figures 1c & 1d).

**Figure 2.**
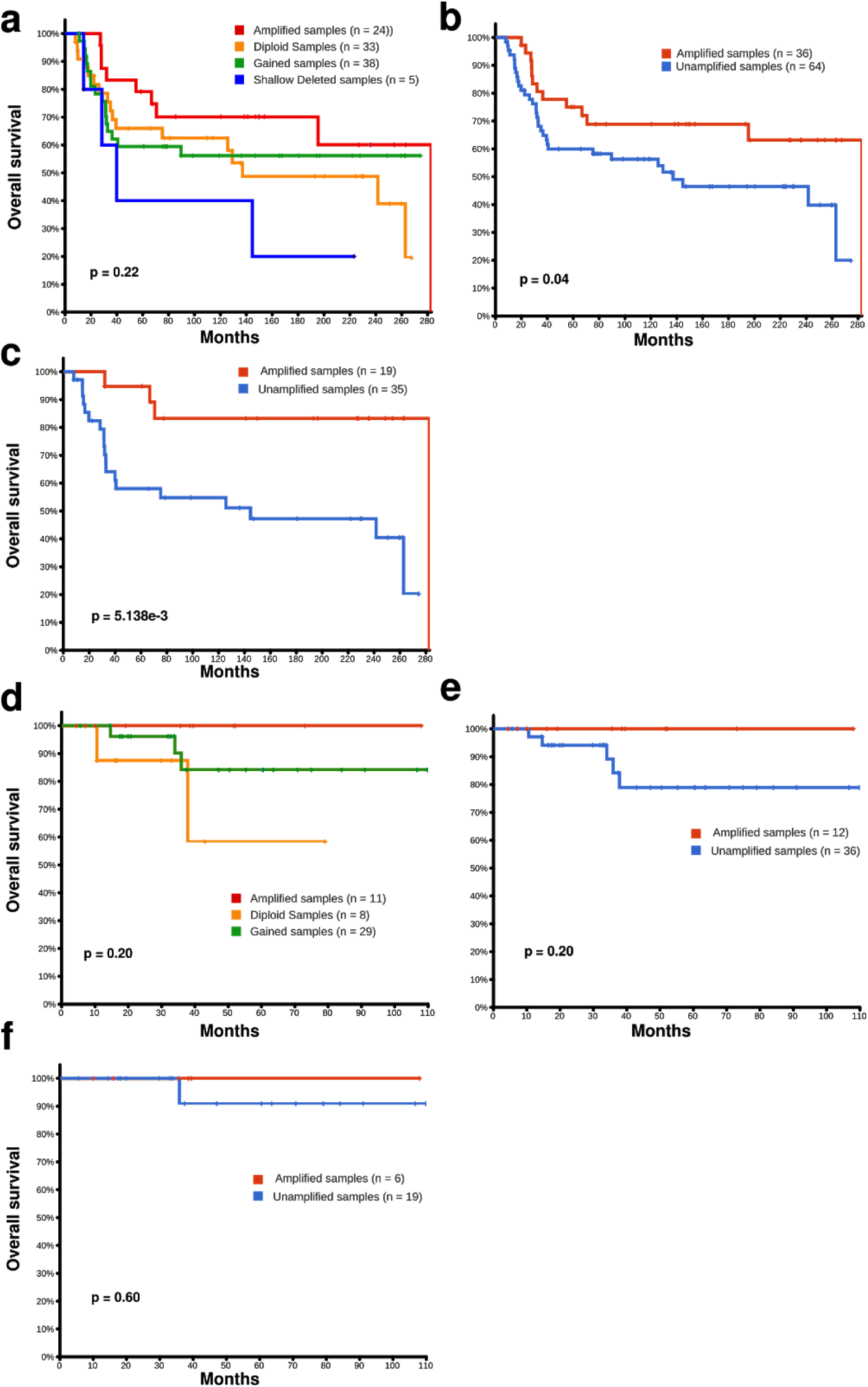
a, b & c) Kaplan-meier overall survival curves of treated HR-/HER-METABRIC cohort (n = 100) stratified by their copy number status (subfigures a & b) and/or stage (subfigure c). d, e & f) Kaplan Meier overall survival curves of TCGA-BRCa HR-/HER-cohort (n = 48) that are treated post-resection and are stratified by copy number status (subfigures d & e) and/or stage (subfigure f).

To further ascertain the role of this locus alteration in disease progression and prognosis, we evaluated the distribution of positive lymph node counts stratified by CN status across the treated samples using the METABRIC dataset. We observed that amplified samples have considerably lower lymph node involvement (p = 0.053) relative to unamplified samples (figure 1e).

Amplified samples over-expressed cyclin proteins (CCNB1, CCNE), cyclin dependent-kinases (CDK1, CDK6 & CDK8), mismatch repair proteins (MSH2 and MSH6), minichromosomal maintenance protein complex (MCM) genes and other markers of DNA replication and cell proliferation. Moreover, TCGA TNBC amplified samples over-expressed cell cycle proteins (CHEK1, RAD50, FOXM1). When using the gene expression data of TNBC patients (GEO accession: GSE25066) who either achieved pCR or relapsed (n = 178), we observed similar upregulation of mitotic pathways, cell proliferation, and chromosomal organization pathways in the pCR cohort (figure 3a). Further, using GSE110153 we observed up-regulation of mitotic pathways in docetaxel sensitive xenografts models relative to the resistant models. We also noticed over-expression of numerous 1q genes in sensitive models, suggesting 1q copy number gain/amplification (figure 3b).

**Figure 3.**
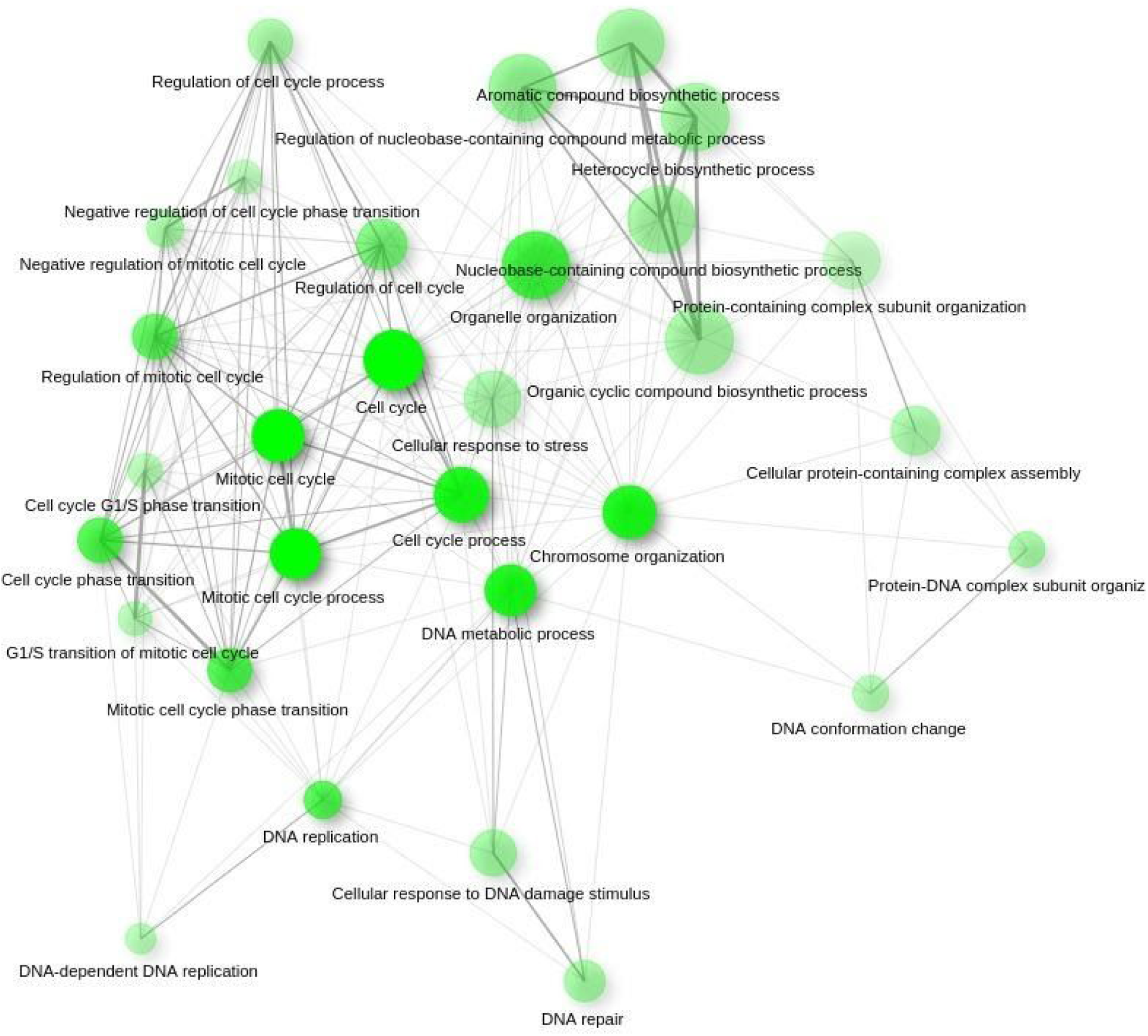
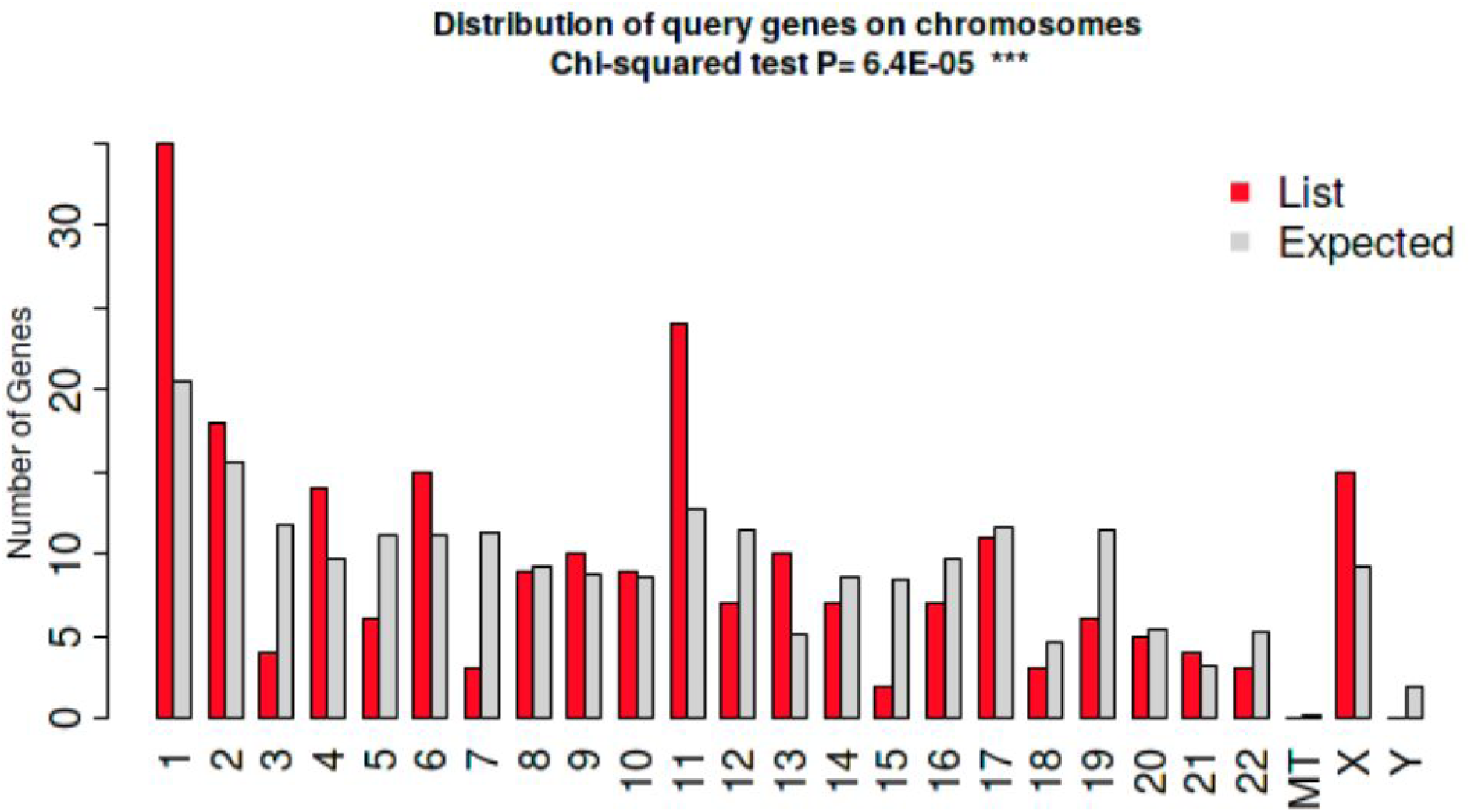
a) Network plot illustrating upregulated GO biological processes in pCR achieved cohort (GSE25066). The circumference of the node represents the size of the gene sets. The color intensity of the node represents their enrichment significance. The thickness of the edges represents the extent of the overlapping gene sets. b) Chromosomal representation of significantly overexpressed genes (p < 0.05) in Docetaxel Sensitive Xenograft models (GSE110153).

### 1q21.3 amplified tumors show differential immune cell enrichment

We used the CIBERSORT to perform immunophenotyping using METABRIC RNA-seq data on samples stratified by both CN and treatment status (16). We observed several differences in the immunological landscape across the cohorts. In particular, higher monocyte and M1 macrophage content and lower M2 macrophage content were observed in treated, amplified samples (figure 4a).

**Figure 4.**
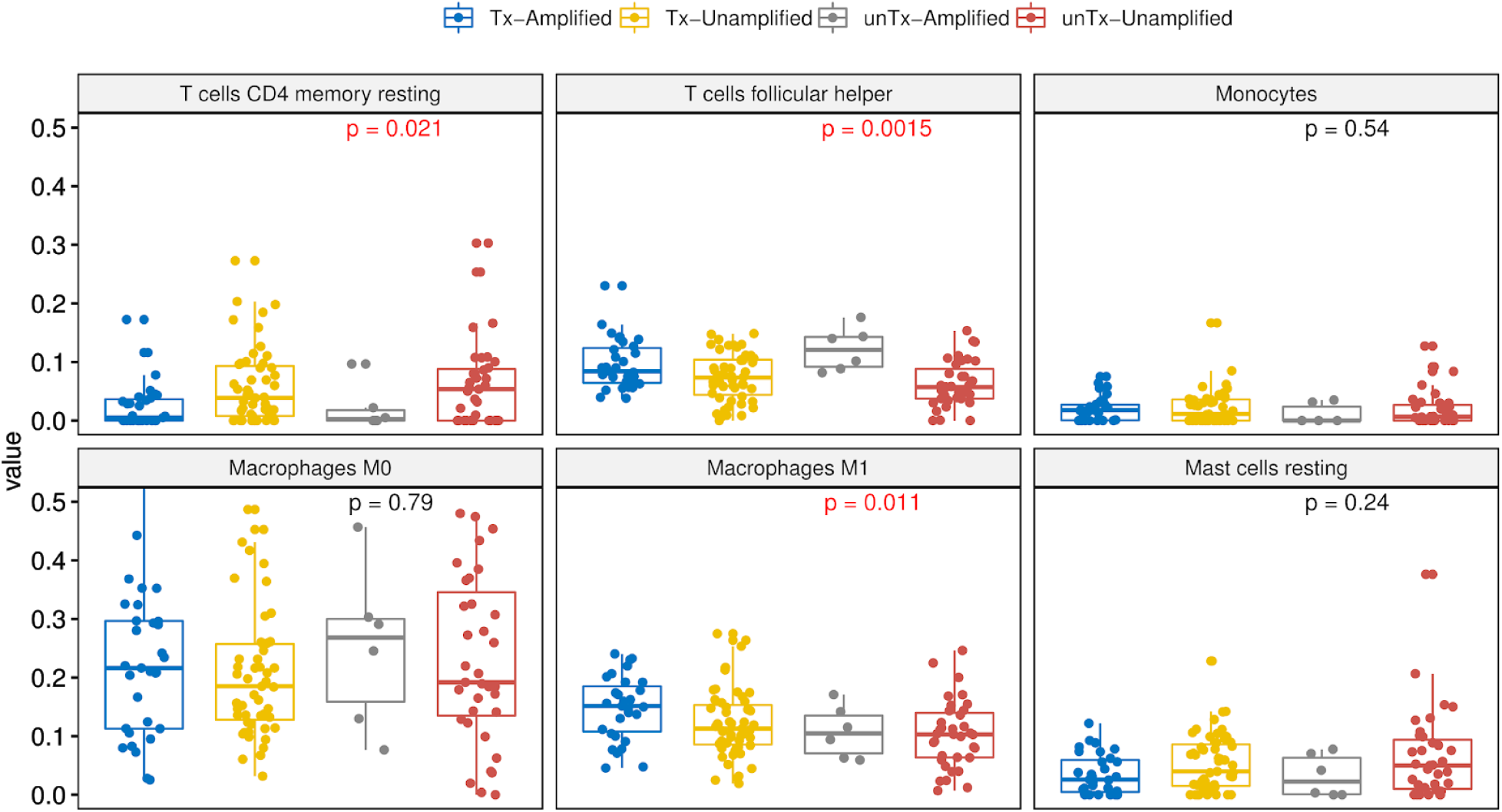
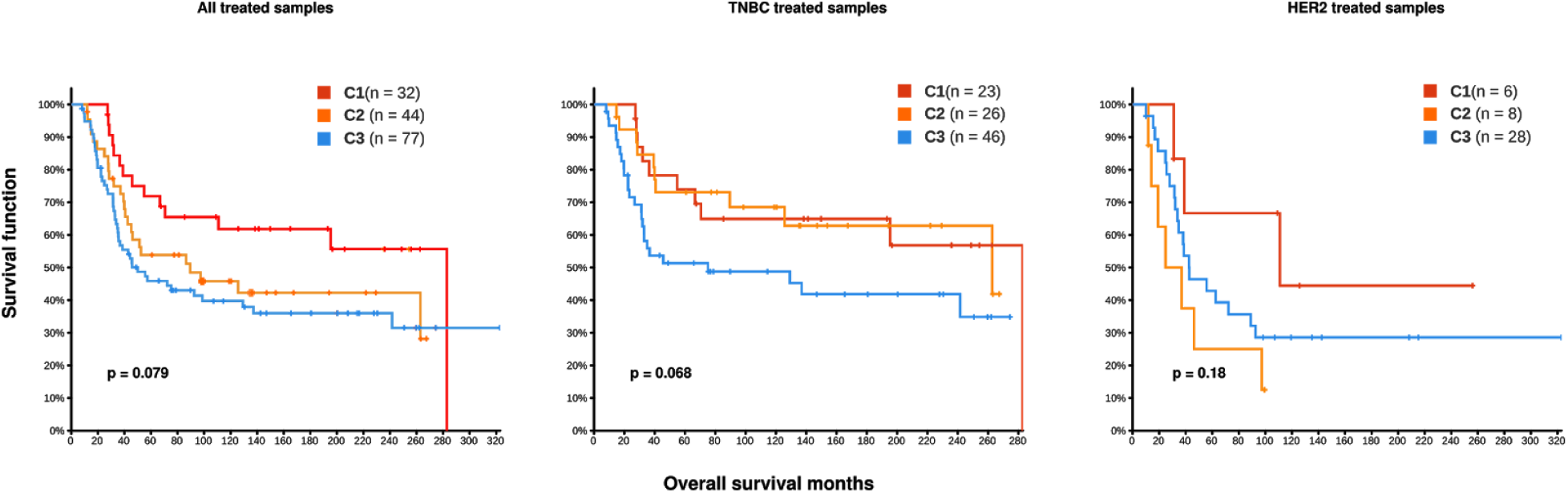
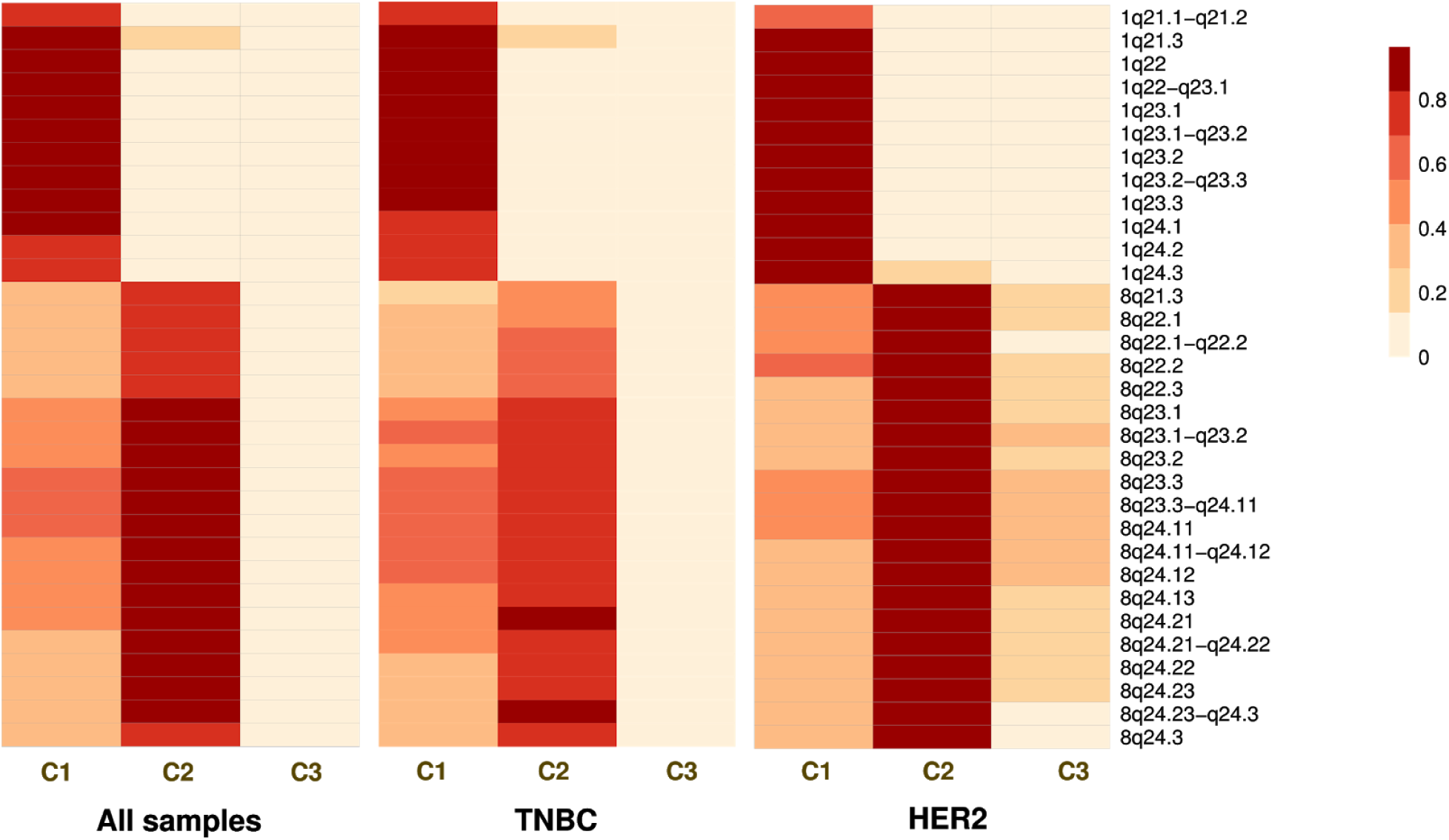
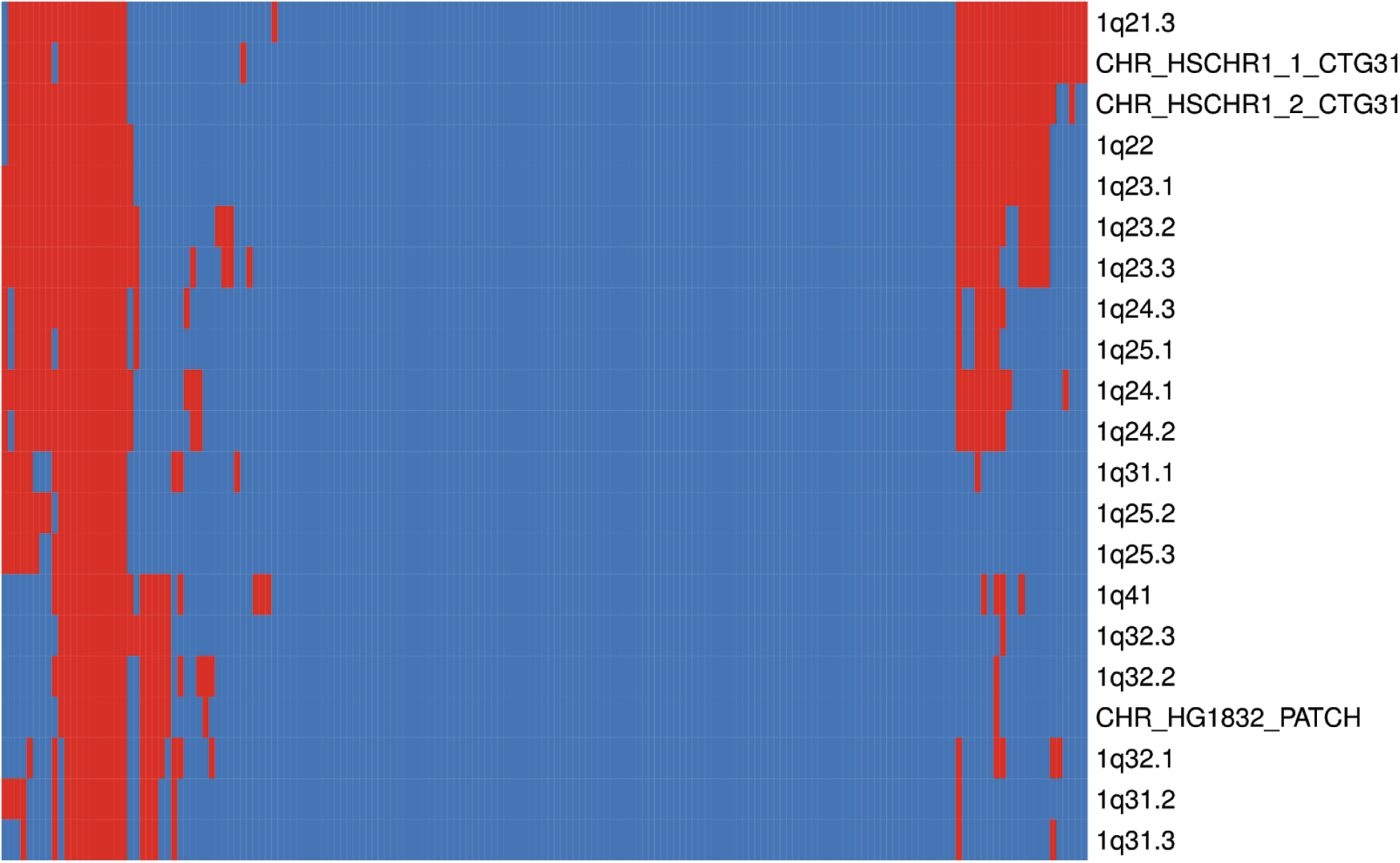
a) Boxplot illustration of immune cell content across METABRIC TNBC cohort grouped with respect to 1q amplification and treatment status. Tx-Amplified = 1q21.3 amplification, treated samples (n = 31); Tx-Unamplified = No 1q21.3 amplification, treated samples (n = 51); unTx-Amplified = 1q21.3 amplification, untreated (n = 6); unTx-Unamplified = No 1q21.3 amplification, untreated (n = 35). Pretreated amplified samples show higher expression of monocytes and M1 macrophages. In addition, amplified samples irrespective of treatment show higher expression of tfh signatures. b) Survival curves of sample clusters (C1, C2 & C3) of three independent analyses (all, TNBC and HER2 treated cohorts) using hierarchical clustering approach on METABRIC gene-level copy number data. Log rank p values are reported. c) Schematic representation of amplification incidence of 1q and 8q loci across all (n = 173), TNBC only (n = 100) and HER2 only (n = 42) treated samples. Differential analysis of the cohorts were done using the cbioportal custom analysis feature. Significant gene level amplification incidence rates (q < 0.05) of 1q and 8q genes are transformed to locus-level incidence using the median incidence across the genes of their corresponding locus. Only loci with incidence proportion of > 0.70 in the clusters were included for the heatmap. d) Clustered heatmap of significantly co-occurring loci across 173 treated METABRIC TNBC BRCa samples.

Although M1 macrophage infiltration was shown to promote anti-tumor activity, a recent study of trastuzumab treated metastatic HER2+ breast cancer tumors demonstrated that higher infiltration of M1 macrophages and CD8+ T-cells is significantly associated with improved overall survival, with longer progression free state after response to treatment (17,18). To confirm this finding, we performed immunophenotyping using gene expression data of matched pre-chemotherapy and post-chemotherapy breast cancer samples (NCBI GEO accession: GSE28844) that were either treated with anthracyclines or taxanes. We observed higher fractions of monocytes in post-chemotherapy samples and M1 macrophages in pre-chemotherapy samples with higher Miller and Payne grades (grades 4 and 5).

In previous studies, it was shown that chemotherapy associated elevations in blood monocytes confers a survival advantage (19). However, we propose that increases in monocytes with higher infiltration of M1 macrophages during/after chemotherapy can elicit robust anti-tumor responses.

Irrespective of treatment status, amplified samples demonstrated significantly higher infiltration of T-follicular helper cells (Tfh). Tfh cells are shown to intensify the humoral response and to promote anti-tumor activity. Especially in aggressive breast cancer subtypes like TNBC and HER2+/HR-, expression of Tfh signatures shows an almost linear relationship with survival (20).

### Unsupervised clustering reveals the influence of 8q amplification on survival

A considerable fraction (40-60%) of the METABRIC and TCGA TNBC samples with 1q21.3 amplification also possessed concurrent amplification events in 8q loci. These samples with concurrent amplification events also over-expressed mitotic and cell cycle regulatory genes indicating the retention of mitogenic function.

We used unsupervised hierarchical clustering to group all METABRIC chemo-radiotherapy treated samples (n = 173) based on all gene-level copy number alterations. As expected, 83% of all the treated samples were either HR-/HER+ or TNBC. Three main clusters were obtained [C1 (n = 32), C2 (n = 44) & C3 (n = 77)] where considerable survival differences among these clusters were observed with a median survival of 132, 69 and 45.63 months respectively (figure 4b). There were no significant differences in stage distribution, cellularity and age across the groups. Almost all samples in the C1 cluster were ER-(Table 1). Approximately 70% of C1 cluster samples were basal subtype compared to 45% in C2 and 25% in C3. 87% of the C1 samples possessed TP53 mutations compared to 82% of C2 and 66% of C3. All the samples in the C1 cluster possessed 1q21.3 amplification, while a negligible (<7%) number of C2 and C3 samples had these alterations. 8q alterations were predominantly observed in the C2 group (~70%) and to a lesser extent in the C1 group (~45%) (figure 4c). We observed that a considerable fraction (~50%) of the C1 samples also bore 8q23 and 8q24 amplification events in addition to 1q21.3 amplification (q < 0.05). The enrichment of over-expressed genes in the C1 cluster showed up-regulation of mitotic pathways like activation of NIMA kinases and polo-kinase mediated events.

**Table 1.** Histological subtype information of all treated METABRIC samples in 3 main clusters (C1, C2 & C3) yielded by hierarchical clustering

| Cluster | NA | ER-/HER2- | ER+/HER2- High Prolif | ER+/HER2- Low Prolif | HER2+ |
| --- | --- | --- | --- | --- | --- |
| **C1** | 6 | 20 (62.5%) | 1 (3%) | 0 (0%) | 5 (16%) |
| **C2** | 8 | 27 (54%) | 3 (6%) | 1 (2%) | 11 (22%) |
| **C3** | 14 | 45 (50%) | 5 (5%) | 2 (2.2%) | 23 (25%) |

Similar clustering patterns and results were observed when analyzing TNBC (n = 100) and HER2+ (n = 42) samples separately (figure 4c). The TNBC clusters C1 and C2 showed higher incidence of 1q21.3 (~100% and ~25% respectively; q < 0.05) and 8q (~65% and 80% respectively; q < 0.05) amplification incidence compared to C3. Both C1 and C2 showed better survival compared to C3 (p = 0.068) (figure 4b). The genomic and clinical characteristics of the clusters are provided in Table 2.

**Table 2.** Clinical and genetic characteristics of all, TNBC and HER2 METABRIC clusters yielded by hierarchical clustering

|  | Sample size | Median survival (months) | Deceased due to the disease (%) | TP53 mutation (%) | PIK3CA mutation (%) | 1q21.3 amplification frequency (%) |
| --- | --- | --- | --- | --- | --- | --- |
| **All** |  |  |  |  |  |  |
| C1 | 32 | 132 | 34 | 87.5 | 6.5 | 96 |
| C2 | 44 | 69 | 54.5 | 81.8 | 34 | 16 |
| C3 | 77 | 45.6 | 58.4 | 66.2 | 18 | 2 |
| **TNBC** |  |  |  |  |  |  |
| C1 | 27 | 141 | 29.6 | 85 | 3.7 | 96 |
| C2 | 31 | 99 | 35.4 | 93.5 | 9.6 | 19 |
| C3 | 51 | 43 | 51 | 72.5 | 13.7 | 3 |
| **HER2** |  |  |  |  |  |  |
| C1 | 6 | 110 | 33.3 | 100 | 0 | 100 |
| C2 | 8 | 31 | 87.5 | 87.5 | 75 | 12 |
| C3 | 28 | 42.5 | 68 | 64 | 28.5 | 0 |

### Concurrence of amplification events between 1q21.3 and neighboring loci

To evaluate any concurrence of amplification events between 1q21.3 and other (1q or 8q) loci, we performed co-occurrence analysis on chromosomal locus level alterations of 173 treated patients (without regard to histological/molecular subtype). We observed a significant concurrence of amplification events in 1q21.3 adjacent loci (adj-p < 0.05) (figure 4d). None of the 8q loci seemed to show significant amplification concurrence with 1q21.3 locus after p-value correction. These results were reproducibly observed when using the entire METABRIC cohort (n = 2105), irrespective of treatment and subtype status.

### 1q21.3 amplified breast cancer cell line models show higher sensitivity to anthracyclines

We used NCI60 data of all five primary BRCa cell lines to assess the correlation between copy number values of the 1q21.3, 8q23 & 8q24 representative genes and -log10GI50 values of all screened compounds in the corresponding samples. Strong correlations (rho > 0.70; p < 0.10) were observed between 1q21.3 copy number values and -log10GI50 of 46 compounds (figure 5a). In addition, we performed principal component analysis on the correlations between-log10GI50 values of 11 cytotoxic drugs and copy number values of the representative genes, and observed strong associations between 1q CN values and inhibitory concentrations (−log10GI50) of fluorouracil, gemcitabine, anthracyclines, alkaloids, fulvestrant and chlorambucil. (figure 5b).

**Figure 5.**
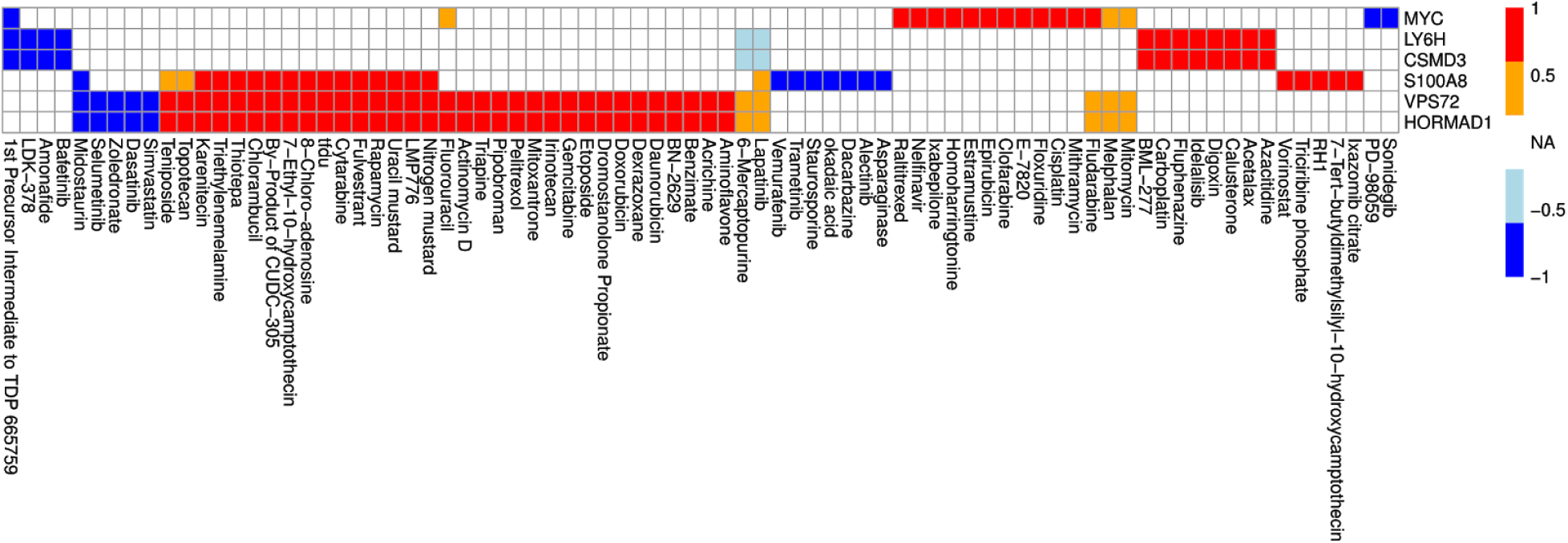
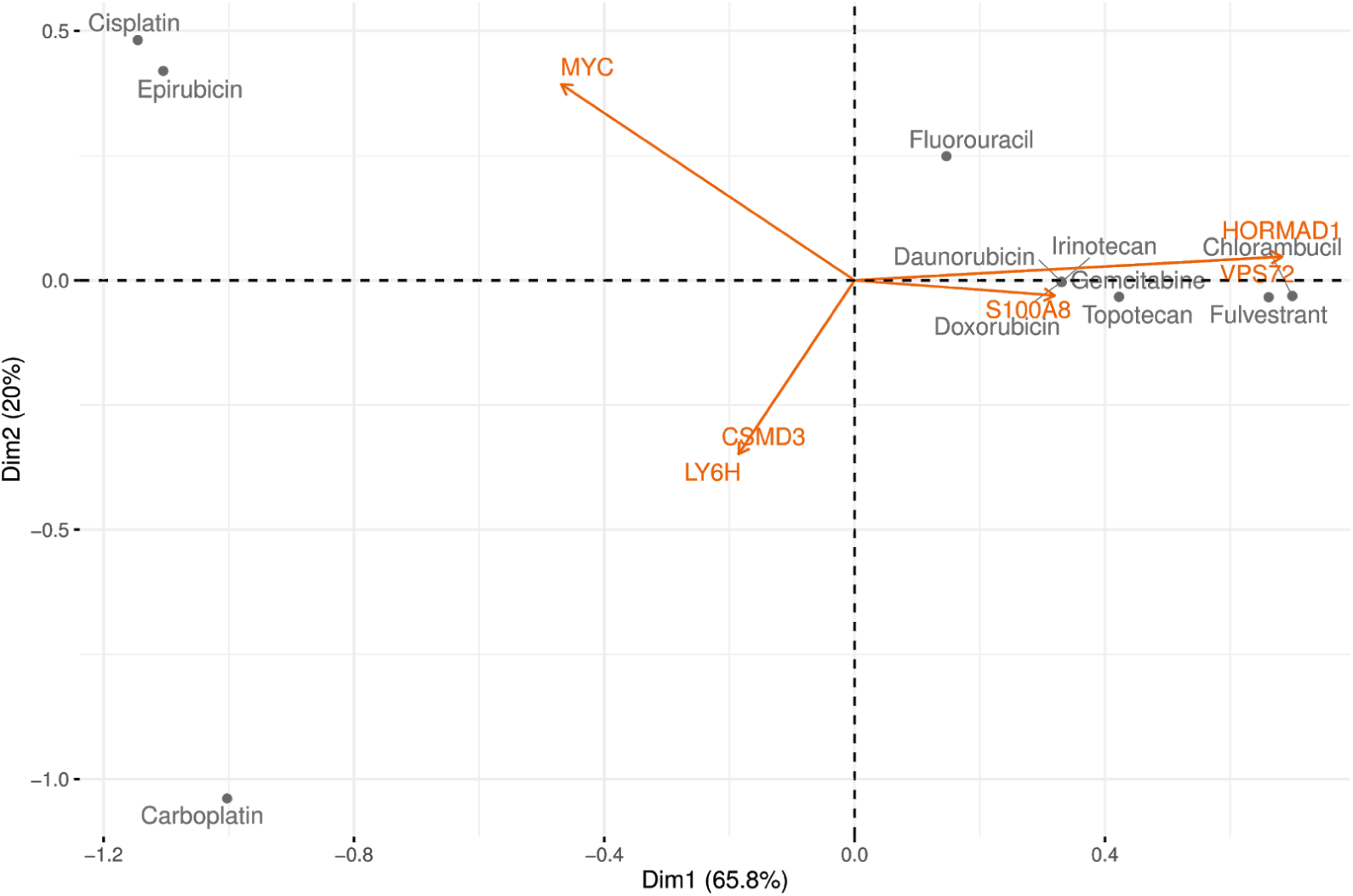
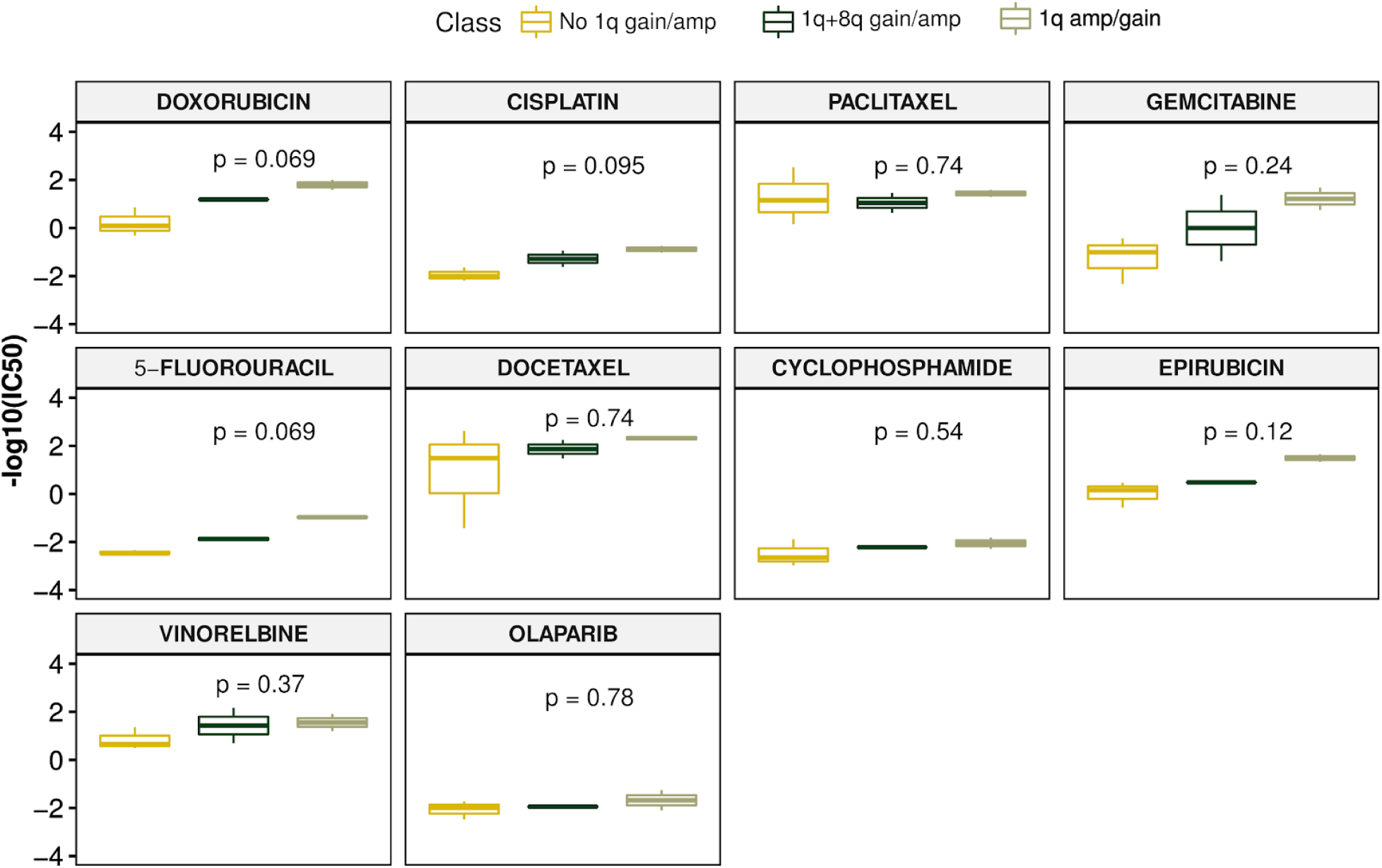
a) Heatmap depicting the strength of correlation (p < 0.10) between -logGI50 values of compounds and copy number values of 1q21.3 and 8q representative genes (NCI60 data). b) Principal component plot illustrating the association between 1q and 8q copy number status and 11 chemotherapeutic agents (NCI60 data). The spearman correlation values between screened compounds and copy number values (p < 0.10) are used to perform principal component analysis. Strong associations between 1q21.3 copy number status and sensitivity of fluorouracil, gemcitabine, anthracyclines, alkaloids, fulvestrant and chlorambucil can be discerned. c) Boxplots of drug sensitivity profiles (−log10IC50) of cell lines stratified by 1q and 8q copy number status across 10 breast cancer drugs are depicted (CCLE & GDSC data). 1q21.3 amplified cell lines showed significantly high sensitivity (p < 0.05) for doxorubicin, cisplatin and 5-fluorouracil.

Using the Cancer Cell Line Encyclopedia (CCLE) data, we identified that the DU4475 TNBC cell line possessed 1q21.3 amplification. Many comparative studies have already demonstrated higher chemotherapeutic responses of this cell line (21,22). We used CCLE 2019 copy number data to identify and stratify the ER-/HER-cell lines with respect to 1q and 8q copy number statuses. Further, we used Genomics of Drug Sensitivity in Cancer (GDSC) dataset to ascertain the sensitivities of 1q gain/amplification (n =2), 1q+8q gain/amplification (n=2) and without 1q amplification TNBC cell lines (n = 3) across ten cytotoxic drugs (Table 3). Although we noticed high sensitivity of 1q amplified/gained cell lines for most of the drugs, significant differences (p < 0.05) in sensitivity were observed with doxorubicin, cisplatin, and 5-fluorouracil (figure 5c).

**Table 3.** The presence of RB1 homozygous deletion, PIK3CA and TP53 mutations across TNBC cell lines classified according to the presence of 1q or/and 8q gain/amplification for drug sensitivity analysis.

|  | **TP53**<br>**mutation** | **PIK3CA**<br>**mutation** | **RB homozygous deletion** |
| --- | --- | --- | --- |
| **No 1q gain/amplification** |  |  |  |
| HS578T | **X** |  |  |
| CAL120 | **X** |  |  |
| HCC1937 | **X** |  | **X** |
| **1q + 8q gain/amplification** |  |  |  |
| HCC1599 | **X** |  |  |
| HCC2157 | **X** |  | **X** |
| **1q gain/amplification** |  |  |  |
| DU4475 |  |  | **X** |
| CAL148 | **X** | **X** | **X** |

### Chemo-radiotherapy treated samples have higher incidence of 1q21.3 amplification

We observed a two-fold increase in the 1q21.3 amplification incidence in treated (36%; n = 100) relative to untreated METABRIC TNBC samples (17%; n = 48). We hypothesize that this difference in amplification rates may be due to treatment induced chromosomal breaks and its association with resulting amplicons. However, it should be noted that no differences in amplification rates were observed between TCGA samples with or without prior cancers. Moreover, we used the MSKCC 2017 targeted sequencing BRCa data to evaluate the incidence of amplification events across genes belonging to loci 1q21.2, 1q22 & 1q23.1 between primary (n = 481) and metastatic tumors (n = 783) and no significant differences in the amplification rates were observed across metastatic and non-metastatic primary tumors. We used genes in 1q21.3 adjacent loci as none of the genes in the 1q21.3 locus were profiled for copy number alterations in this study and also our earlier analysis showed significant concurrence of amplification events between the 1q21.3 locus and the neighboring loci.

Although there are isolated studies showing evidence of a strong association between chemotherapy and genomic instability, only a few studies highlight the association between chromosomal instability, 1q21.3 alterations, and therapy response (23). To determine whether chromosomal breaks follow a non-random pattern, we used HumCFS database to identify most common chromosomal fragile sites (24). We observed that the highest number of fragile sites are present on Chromosome 1 and 2 (13 sites each). In particular, the FRA1F fragile site induced by aphidicolin, an inhibitor of DNA replication, spans most of the 1q21 region (including 1q21.3 locus).

It is already known that aphidicolin acts synergistically to improve the sensitivity and cytotoxic potential of purine analogs and platinum compounds (25). It is also established that entinostat, a HDAC inhibitor, when combined with low-dose doxorubicin with or without All Trans Retinoic Acid (ATRA) decreases cell viability and tumor volume, in addition to inducing cell differentiation of tumor initiating cells in cell lines and xenograft models (26). Using Gene Expression Omnibus (GEO) dataset GSE63351, we observed increased representation (compared to expected probability) of chr 1q genes in over-expressed gene set in entinostat (+/-ATRA) and low-dose doxorubicin combination treated MDA-MB-231 cells when compared to low-dose doxorubicin treated MDA-231 cells, suggesting a gain or amplification of chr 1q (figure 6). The MDA-MB-231 is an aggressive, highly invasive metastatic TNBC cell line that does not possess any alterations in the 1q region at the baseline, and it is only partially responsive to doxorubicin.

**Figure 6.**
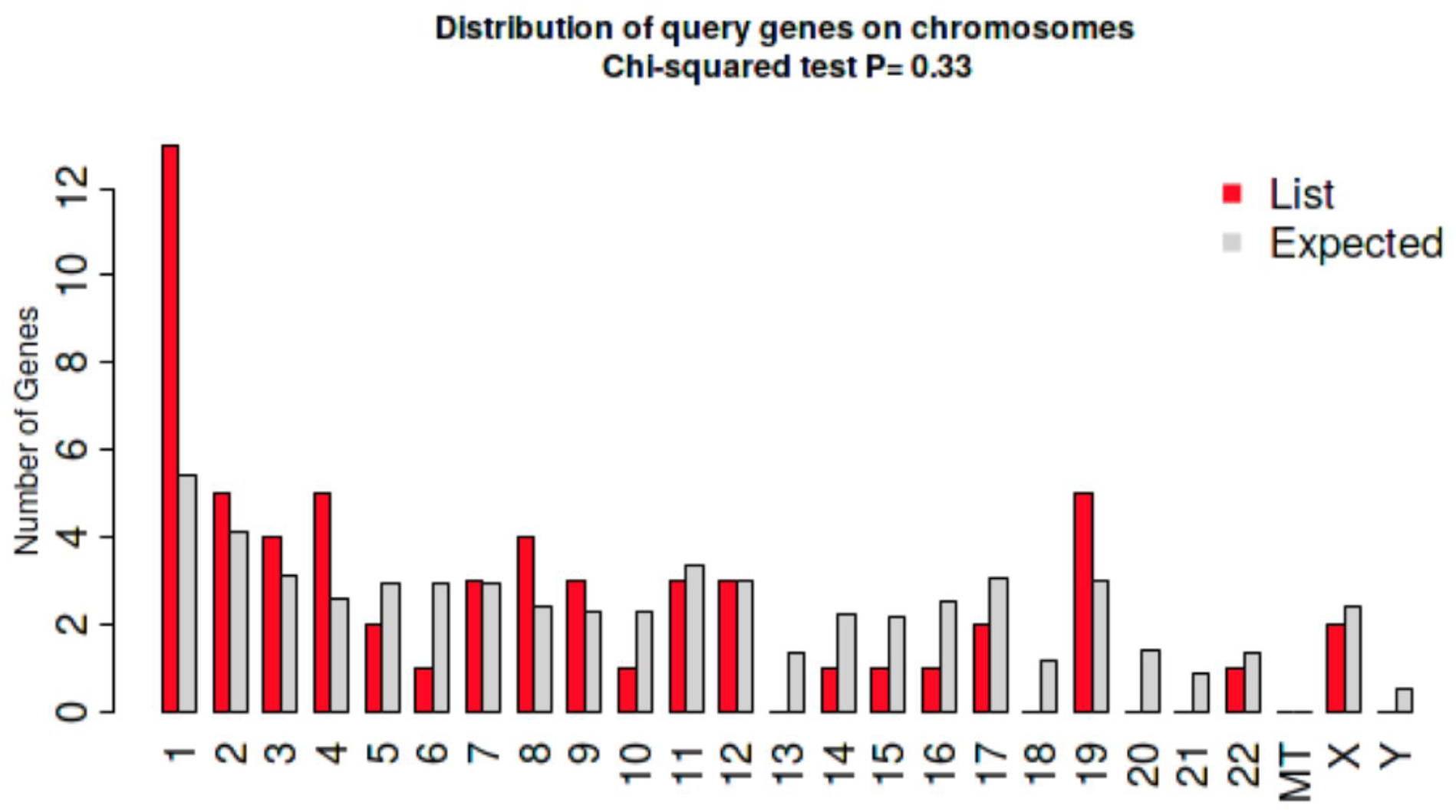
Chromosomal representation of significantly overexpressed genes (Adjp < 0.05) in Doxorubicin + Entinostat treated group compared to Doxorubicin treated group in MDA-MB-231 cells.

### 1q21.3 amplified tumors share similarities across cancer types

Basal subtype BRCa shares similar molecular features with serous uterine carcinomas (UCEC) and high grade serous ovarian carcinomas (HGSOV) (27). Therefore, we evaluated whether our findings could also apply to TCGA Uterine Endometrial Carcinoma (UCEC) data. We observed that 95% (37/39) UCEC samples with 1q21.3 amplification belonged to the “CN High” subtype. Most of the “CN high” subtype samples have shared membership with “Mitotic” mRNA subtype and are characterized by highest transcriptional activity in addition to increased dysregulation of the cell cycle pathways (27). Serous subtype cancers of OV and UCEC are considered most aggressive (28). Majority of the serous subtype samples of TCGA UCEC cohort (97 of 109 samples) are also classified as “CN high” tumors and these patients are shown to have poor survival outcomes (27). However, we noticed that 25% of TCGA-UCEC serous subtype samples that bore 1q21.3 amplification have better disease free survival (median survival of 27 months vs 21 months) (p = 0.08) with fewer event rates (4/14 (28.5%) vs 26/49 (53%)) compared to unamplified tumors. We also observed high incidence rates (~12%) of 1q21.3 amplification across TCGA HGSOV tumors. A subset of these patients (n = 30) with 1q21.3 but not 8q amplification events demonstrated better overall and disease-free survival relative to other patients (median survival of 45.5 months vs 33.54 months) (log rank p value = 0.12).

## Discussion

It is long known that genotypic characteristics of the cell inherently possess the potential to modulate the therapy response. Our results exhibit that 1q21.3 amplified tumors possess increased cell cycle dysregulation alongside apparent mitogenic phenotype and thus show improved sensitivity to cytotoxic therapies. We also observed intensified pro-inflammatory tumor response in these tumors.

Over-expression of mitogens, cyclins and cell cycle regulators indicate the high proliferative potential of untreated 1q21.3 amplified tumors. In particular, the increased expression of FOXM1, CDK, CCNB1 and CCNE proteins in these samples further points to the high mitotic index of these tumors. In treated samples, however, high mRNA expression of AURORA and BUB1 suggests mitotic checkpoint arrest. Mitotic catastrophe is a well-established phenomenon whereby a cell—when exposed to anti-cancer treatments—undergoes apoptotic cell death after aberrant mitosis. Improved sensitivity of cytotoxic agents towards amplified tumors may be due to enhanced DNA fragmentation ability.

Our results exhibit the contrariety of 1q21.3 amplified tumors: chemo-radiosensitivity and high aggressive potential. We hypothesize that amplified tumors demonstrate better histological response to standard cytotoxic agents and likely achieve remission. However, in case of late diagnosis or deferred treatment, the risk of rapid progression and invasiveness may be very high.

Based on our cell line drug sensitivity results, we assume that 1q21.3 amplified tumors are more responsive to anthracyclines. It is shown in earlier studies that neoadjuvant anthracyclines are more effective in basal and HER2+ subtypes for achieving pCR (29). Interestingly, these subtypes have the highest incidence of 1q21.3 amplification events and therefore, we postulate that patients with amplified tumors are most likely to achieve pCR with neoadjuvant/preoperative chemotherapy. Our analysis involving tumors/cell lines without amplification events hint higher risk of treatment refractoriness with the use of anthracyclines. Taxanes may be useful in the optimal management of these tumors due to their negligible differences in their sensitivity across the amplified and unamplified cell lines. Moreover, in current clinical practice, taxanes are generally preferred over anthracyclines in the treatment of refractory and advanced cases (30). Our results also indicate that HDAC inhibitor (HDACi) entinostat like its cousin vorinostat, facilitates chromosomal breaks while preventing DNA repair and thus sensitizing tumor cells to cytotoxic agents (31). This action likely enhances the susceptibility of unamplified tumors to DNA-damaging agents. However, their potential use in the treatment of refractory tumors is yet to be ascertained.

Although other BRCa subtypes like luminal cancers showed high incident rates of 1q21.3 amplification, there were no considerable prognostic differences observed with respect to the 1q21.3 CN status across these subtypes. On the contrary, the shared prognostic and molecular features of 1q21.3 amplified tumors across TNBC, HGSOV and serous UCEC suggest the possibility of a unified treatment approach.

## Methods

### Data retrieval

We retrieved TCGA and METABRIC RSEM normalized RNA-seq data (FPKM-UQ), normalized RPPA protein expression data, GISTIC 2.0 gene-level copy number data and clinical information from cbioportal data repository. GSE28844, GSE25066, GSE110153 and GSE63351 normalized data were obtained from NCBI Gene Expression Omnibus (GEO) repository.

NCI60 copy number and drug sensitivity data was downloaded from UCSC XENA (https://xena.ucsc.edu/public/). CCLE copy number data and Genomics of Drug Sensitivity in Cancer (GDSC) 1 & 2 datasets were obtained from DepMap webportal (https://depmap.org/portal/).

### TCGA and METABRIC BRCa analyses

Only primary tumors were used for the analysis. Samples with missing clinical information regarding HR/HER receptor status, treatment status, sample type and neoadjuvant therapy status were discarded prior to the stratification. Only treatment naive TCGA TNBC primary samples were considered for the analysis. TNBC samples were identified by HR and HER receptor negativity provided in the clinical information.

Unless specified, the classification of copy number alterations are based on widely accepted copy number value thresholds where deep/homozygous deletion (−2), arm-level/shallow/heterozygous deletion (−1), diploid (0), arm-level/gain (1) and amplification (2). We focused most of our analyses on amplification (focal/high level gain of gene copies) rather than copy number gain as their contribution can be determined clearly. Nevertheless, we used arm-level alterations sporadically to evaluate the consistency of the results.

We used cbioportal (www.cbioportal.org) to retrieve significantly over-expressed genes (P < 0.05) in samples with amplification events in 1q21.3 representative genes (S100A8, HORMAD1 & VPS72) across TNBC samples of TCGA-BRCA (n = 112), METABRIC treated cohort (n = 100) and METABRIC untreated cohort (n = 48). Survival analysis was performed using samples treated with chemo-radiotherapy (METABRIC (n = 100) & TCGA-BRCA (n = 48)). We used clinical information on treatment status of TCGA-BRCA samples from Genomic Data Commons (gdc.cancer.gov) and receptor (HR & HER) status information from TCGA-BRCA firehose legacy dataset. Consensus (intersection) of copy number statuses of representative genes was used for survival analyses of samples grouped by specific copy number status (Amplification, Diploid, Gain and Shallow deletion). Harmonized stage-specific survival analysis was performed using AJCC Stage II samples (METABRIC (n = 57) & TCGA-BRCA (n = 25)). Log-rank test p-values are reported for the survival curves.

Immunophenotyping was performed on previously normalized METABRIC data using CIBERSORT tool with LM22 signatures and 500 permutations. Quantile normalization was disabled for all the analyses. We only considered samples with CIBERSORT p-value < 0.05 for further analysis. We profiled the immune cell content across samples grouped by treatment and 1q amplification status. Kruskal Wallis test (p < 0.05) was used to evaluate the significance of differential immune cell enrichment.

The *ggplot2*, *ggpubr, survminer and factoextra* R packages were used to make boxplots, violin plots and barplots. Survival plots were made using cbioportal’s in-built custom analysis feature. Network and chromosome distribution plots of differential genes were made using ShinyGO web browser (32). GO based enrichment analysis using GO biological processes was performed using FUNSET web browser (33).

### Co-occurrence analysis

Pairwise co-occurrence analysis on METABRIC 1q and 8q copy number data of treated samples (irrespective of the histological subtype) (n = 173) was carried out using the *DISCOVER* python package (34). The gene-level copy number alterations were transformed to locus-level alteration by using the median copy number values across the locus. Arm-level copy number alterations were not considered for this analysis. An independent analysis of similar scope was also performed using the entire METABRIC cohort (n = 2105). FDR corrected p-values (q value < 0.05) of pairwise comparisons were used to infer concurrence of amplification events.

### Hierarchical clustering analysis

We used the gene-level processed copy number values of all 173 chemo-radiotherapy treated METABRIC samples (irrespective of their histological subtype) for the analysis. Hierarchical clustering was performed using *hclust* R base function with complete linkage. For all the three analyses involving all, TNBC and HER2 treated samples, we used the root node to separate the C1 cluster from C2 and C3. We seperated C2 and C3 using the “all” treated cohort by making a cut at level 3 of the tree. We made a cut at level 4 to separate clusters C2 and C3 of “TNBC” clusters. C2 and C3 clusters of HER2 treated samples were obtained with a cut at level 1 of the tree.

### NCI60 and GDSC Cell line analyses

Spearman correlation was performed between gene level copy number values (1q and 8q representative genes) of 5 NCI60 breast cancer cell lines (MCF7, MDA-MB-231, HS 578T, BT-549 & T-47D) and their corresponding drug concentration values that cause 50% growth inhibition (−logGI50) for 262 screened compounds. We used a permissive p-value threshold of 0.10 due to small sample sizes to identify significant correlations. The correlation values with respect to 11 cytotoxic agents were used to perform principal component analysis.

We used the DepMap portal (https://depmap.org/portal/) for identification of ER-/HER-breast cancer cell lines. Using the CCLE cell line gene-level copy number values, we stratified 23 ER-/HER-cell lines into three groups a) 1q amplification/gained, b) 1q+8q gain/amplification and c) No 1q amplification. We used an arbitrary mean copy number value of > 1.5 of 1q and 8q representative genes to identify amplification events across the cell lines.

### Gene expression datasets

We used CIBERSORT tool for immunophenotyping analysis on GSE28844 gene expression data to identify differential immune cell enrichment in pre and post chemotherapy treated samples (n = 56) with higher Miller and Payne grades (grade 4 & 5). We used Kruskal Wallis test (p < 0.05) for assessing the significance of differential enrichment.

To establish the association between up-regulation of mitotic pathways and treatment sensitivity, pathway enrichment using *ShinyGO* (http://bioinformatics.sdstate.edu/go/) was performed on significantly over-expressed genes (adjusted p < 0.05) in pCR achieved cohort of 178 TNBC cohort (NCBI GEO accession: GSE25066). Differential expression analysis was performed using Mann-Whitney U test and p values were corrected using the FDR method. Identification of up and downregulation of significant differentially expressed genes were done by calculating the median across the cohorts.

Differential expression analysis of docetaxel sensitive and resistant xenografts (NCBI GEO accession: GSE110153) was performed using NCBI GEO “GEO2R” functionality. The chromosomal representation analysis and pathway enrichment were performed using *ShinyGO* web server.

To evaluate the effect of HDAC inhibitor entinostat on MDA-MB-231 cells, we used gene expression data of low-dose doxorubicin treated and low-dose doxorubicin and entinostat combination treated MDA-MB-231 cells (NCBI GEO accession: GSE63351). We used “GEO2R” functionality of NCBI GEO to perform differential gene expression analysis. We assessed two p-value thresholds (adjusted p < 0.05 and nominal p < 0.05) and performed chromosomal representation analysis of over-expressed genes in combination treated cells using *ShinyGO* web server.

